# An ultra-low dose of Δ9-tetrahydrocannabinol improves Alzheimer’s Disease-related cognitive deficits

**DOI:** 10.1101/2021.08.22.457252

**Authors:** K. Nitzan, L. Ellenbogen, T. Beniamin, Y. Sarne, R Doron

## Abstract

Alzheimer’s disease (AD) is the most common form of dementia. AD has a physical, emotional, and economic impact on the patients and their families and society at large. More than a decade since its discovery, there is still no available treatment. Δ9-tetrahydrocannabinol (THC) is emerging as a promising therapeutic agent. Using THC in conventional-high doses may have deleterious effects. Therefore, we propose to use an ultra-low dose of THC (ULD-THC). We previously published that a single injection of ULD-THC elevated Sirtuin-1 (Sirt-1) levels in the brain and ameliorated cognitive functioning in several models of brain injuries as well as in naturally aging mice. Our working hypothesis suggests that ULD-THC can prevent and even reverse AD pathology. In this preliminary study, we saw that a single injection of ULD-THC alleviated cognitive impairments of a mice model for AD, 5xFAD mice. Our work may establish the foundations for the development of a pharmaceutical preparation for the treatment of AD patients, thus, bringing the ULD-THC treatment closer to clinical application.

## Introduction

The world’s population is aging. With better health care and sanitation, and with greater availability of food, people worldwide are living longer: according to a UN report from 2017, 12.7% of the world’s population is aged 60 and over, and this number is expected to double in the next 20 years [1]. Aging is associated with many neurodegenerative disorders, among them Alzheimer’s Disease (AD). As AD usually starts to affect people over the age of 60, its incidence has been increasing with the current aging of the population and is becoming a significant public health issue [2]. According to The United States Alzheimer’s Association, 5.8 million Americans aged 65 and older are now living with the disease, and by 2050 their number will double [3]. Recently the FDA approved a new drug for the treatment of AD, Aducanumab, but its efficacy has caused much debate in the scientific community[4, 5]. Thus, although many efforts have been dedicated to finding a cure for AD, there is no effective treatment at present.

AD is a progressive neurodegenerative disease characterized by extracellular senile plaque composed of fibrillar amyloid-β (Aβ) peptides, intracellular neurofibrillary tangles containing hyperphosphorylated tau, brain atrophy, and the disruption of episodic memory function [6]. While much emphasis has been given to the amyloid and tau proteins, AD has a multifactorial etiology. Mitochondrial dysfunction, oxidative stress, unbalanced iron metabolism, and neuroinflammation are only some of the factors involved in the causation of AD [7, 8]. Consequently, treating the end-result of the disease, such as focusing on amyloid beta (Aβ) peptides or tau phosphorylation, has not yielded satisfactory results. What may be needed, instead, is a treatment that involves multiple aspects of the disease.

Cannabinoids are emerging as promising therapeutic agents [9, 10]. Tetrahydrocannabinol (THC) is the major psychoactive constituent of the cannabis plant. More than 565 compounds have been isolated from this plant, of which the two main cannabinoids are Δ9-tetrahydrocannabinol (Δ9-THC) and cannabidiol (CBD) [11]. The endocannabinoid system comprises at least two types of cannabinoid receptors (CBR) – CB1R and CB2R [12], as well as endogenous ligands (e.g., N-arachidonoylethanolamine, AEA and 2-Arachidonoylglycero, 2-AG) and their anabolic and catabolic enzymes. Cannabinoid agonists have been found to be beneficial in a variety of experimental models. In-vitro, under pathological conditions, cannabinoid compounds activation had a beneficial effect on mitochondrial dysfunction [13], and THC demonstrated antioxidative effects [14]. In a cell culture model of Parkinson’s disease, THC had a protective effect mediated through peroxisome proliferator-activated receptor-gamma (PPARγ) activation [15]. In vivo, THC protected against damage from ouabain-induced excitotoxicity [16] and MDMA neurotoxicity [17].

However, one should consider the many limitations of conventional doses of cannabinoids. Numerous studies have shown that THC has deleterious effects [18, 19]. THC exposure in the young has been related to deficits in attention [20], short-term memory [21], and spatial memory [22]. Our previous studies demonstrated the beneficial effects of an ultra-low dose of THC (ULD-THC) in mice. In those experiments, we applied a single extremely low dose of THC (0.002 mg/kg, 1,000-10,000 times lower than the conventional doses used in rodents), which resulted in profound neuroprotective effects. We observed the protective effects of ULD-THC in several models of brain injuries: in a model of epilepsy (PTZ) that impairs cognitive functions [23], in three different CNS insults that cause cognitive deficits in young mice [24]; and, in a mice model of neuroinflammation, in which ULD-THC was protective against cognitive deficits [25]. A single dose of ULD-THC produced a protective effect with a wide therapeutic time window, and the drug could be effectively introduced from 1–7 days before or after the insult. We also observed a beneficial effect in old, senile mice, reflected in a reduction of age-related cognitive decline. [26]. The rationale for using ultra-low doses of THC and the possible mechanisms of action are presented in detail in our recent review [27].

THC treatment may also have the potential to affect many of the factors involved in AD pathologies, such as Aβ clearance, mitochondrial function, inflammation, and neurogenesis [28–30]. Indeed, it was found that, in vitro, THC decreased Aβ concentration in a dose-dependent manner in N2a/APPswe cells [31]. Following our previous findings that a single treatment with an ultra-low dose of Δ9-tetrahydrocannabinol THC (ULD-THC) provided prolonged neuroprotection in response to various brain insults [24, 25], including the improvement of cognitive functions in old, senile mice [26], we performed for the first time a preliminary study to examine ULD-THC as a potential treatment for AD cognitive deficits. Its preliminary data is present in this article.

## Methods

### Animals

All experiments were performed on 1-year old male and female 5xFAD tg-mice (provided by Prof. Arber, The Integrated Cancer Prevention Center, Tel Aviv Sourasky Medical Center, Tel Aviv, Israel) and their wildtype (WT) littermates (n = 8 per treatment group, mixed gender). 5XFAD mice have five mutant human genes associated with AD: Three APP genes, that is, APPsw, APPfl, and APPlon, and two presenilin1 genes, that is, PS1 and PSEN1. Mice were genotyped by PCR analysis of tail DNA to identify the tg-mice, and the non-tg littermates were used as control, WT-mice. Mice were kept in the vivarium at the Ein Kerem Hadassah Medical Center’s Psychobiology Lab of the Open University on a reverse 12 h light/dark cycle and provided with food and water ad libitum. All experiments were performed during the dark phase light (7:00-19:00) under red light. All experimental protocols were examined and approved by the Institutional Animal Care and Use Committee.

### Treatments

#### ULD-THC -

Δ9-THC (donated by Prof. Mechoulam, the Hebrew University, Jerusalem and by NIDA, USA) was dissolved from a stock solution in ethanol into a vehicle solution consisting of 1:1:18 ethanol: cremophor (Sigma-Aldrich): saline, and was administered i.p at a dose of 0.002 mg/kg. Mice were treated with a single i.p. injection of either ULD-THC or vehicle and tested three weeks later.

### Behavioral tests

#### Y maze (YM)

The YM assay is based on the tendency of mice to explore a new environment. The maze comprises three, each 30 cm long and 120° apart identical arms, with each arm presenting different visual cues. The mouse was placed at the distal edge of the start arm during the first session, with one of the two remaining arms blocked and left to explore the maze for 5 minutes. The mouse was then be returned to its home cage and reintroduced into the start arm 2 minutes later for 2 minutes, with the two remaining arms open. The total time that the mouse spent exploring each of the two arms was recorded, normalized, and used to calculate the preference index. [26].

#### Morris’ Water Maze (MWM)

Learning ability and spatial memory were evaluated using MWM. This classical spatial learning assay was performed as previously described by us [26]. Briefly, the mouse was introduced into a round pool at different starting points and allowed 60 seconds to find the platform. Each mouse swam four times per day for three consecutive days. The time and route required for the mouse to find the immersed platform was recorded.

#### Motor function in the Open Field Test

The OFT consists of an empty square arena (40×40×40cm) surrounded by Perspex opaque walls. Mice were placed in the center of the arena, and their behavior was video recorded for a total of 5 min. The maze was thoroughly cleaned with ethanol and allowed to dry between subjects to eliminate any odor cues. Motor behavior was measured by the distance traveled by the mice.

All the behavioral assays were photographed using a video camera, and data were recorded and analyzed using the 13th Noldus EthoVision software.

### Study design

In order to determine the treatment efficacy of ULD-THC in the 5xFAD tg mice model, 1-year-old male and female 5xFAD tg mice and their WT littermates were treated with a single i.p. injection of either ULD-THC or vehicle (n = 8 per treatment group, mixed gender) and tested for short and long-term spatial memory three weeks later.

**Figure 1.**
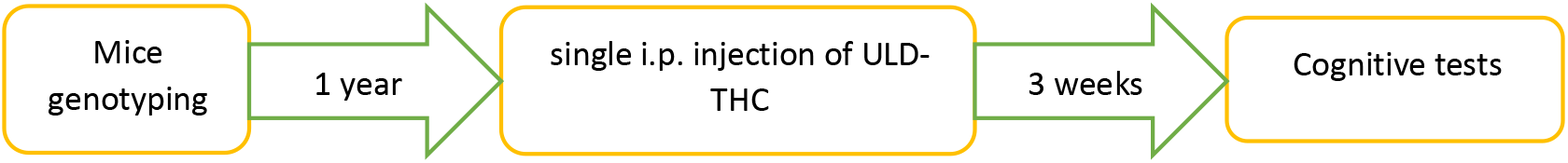
Study design

### Data analysis

All results are presented as mean ± standard error of the mean Behavioral data was analyzed using the ANOVA, further analyzed using post-hoc analysis, or planned contrast. Statistical analysis was performed using IBM SPSS Statistics V.25. The level of significance was set at p <0.05.

## Results

Hippocampus-dependent short-term spatial memory, tested in the Y-maze apparatus (Fig. 2), was impaired in AD-mice compared to WT mice. Importantly, AD-treated mice exhibited a better performance compared to vehicle-treated AD-mice and spent significantly more time in the new arm F(3,28) = 2.975 (p < 0.05).

**Figure 2.**
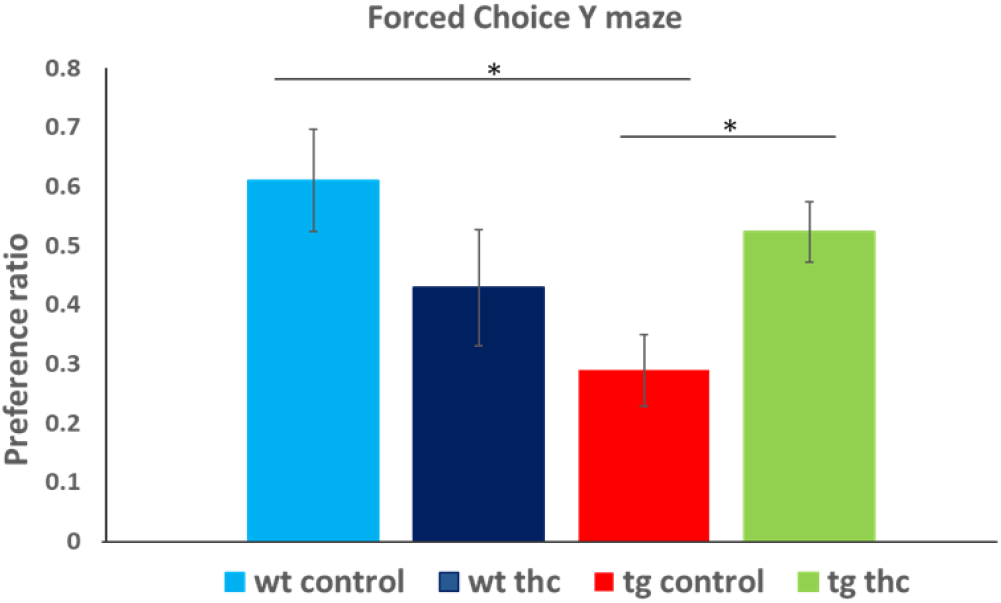
Y maze. one-year-old 5xFad (tg) and control (wt) littermates were treated with a single I.P. injection of ULD-THC or vehicle and tested 3 weeks later (n=8 in each group). A significant difference was found in planned contrast, between the performance of tg and wt mice, with a beneficial effect of ULD-THC treatment in TG mice.

Hippocampus-dependent long-term spatial memory, tested in the Morris Water Maze (was impaired in AD-mice compared to WT mice and the ULD-THC treatment alleviated this impairment. Repeated measures ANOVA for the time to reach the platform yielded significant day*group interaction F(2.5, 60) = 1.735 (p < 0.05). Post-hoc testing revealed on day one a significant difference between the non-treated WT and all other groups. On day two, there was a significant difference between the non-treated 5xFAD and all other groups; on day three, both WT groups performed better than both 5xFAD groups. (Fig 3).

**Figure 3.**
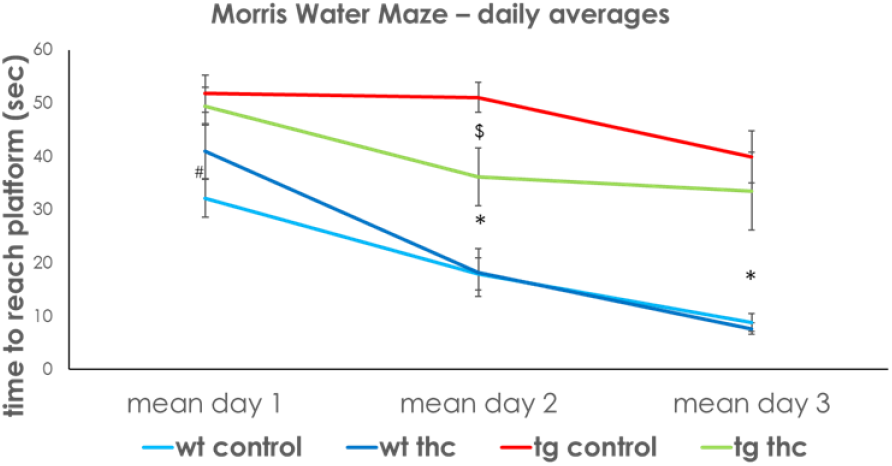
Morris Water Maze. One-year-old 5xFad (tg) and wt littermates were treated with a single I.P. injection of ULD-THC or vehicle and tested 3 weeks later (N=8 in each group). Repeated anova yielded significant interaction. On day 1, post-hoc revealed a significant difference between the non-treated WT and all other group. **On day 2 there was a significant difference between the non-treated tg and all other groups.** On day 3, Both wt groups performed better compared to both tg groups. * Difference between WT groups and TG groups. # Diffrenece between WT control and TG groups. $ Diffrenece between TG treated and TG control groups.

Repeated measures ANOVA for the rout (distance) to reach the platform yielded significant day*group interaction F(5.3, 53.9) = 3.4 (p < 0.05). Pairwise comparisons showed that while both WT mice and treated 5xFad mice significantly improved from day 1 to day three respectively) the non-treated group did not (A). (Fig 4)

**Figure 4.**
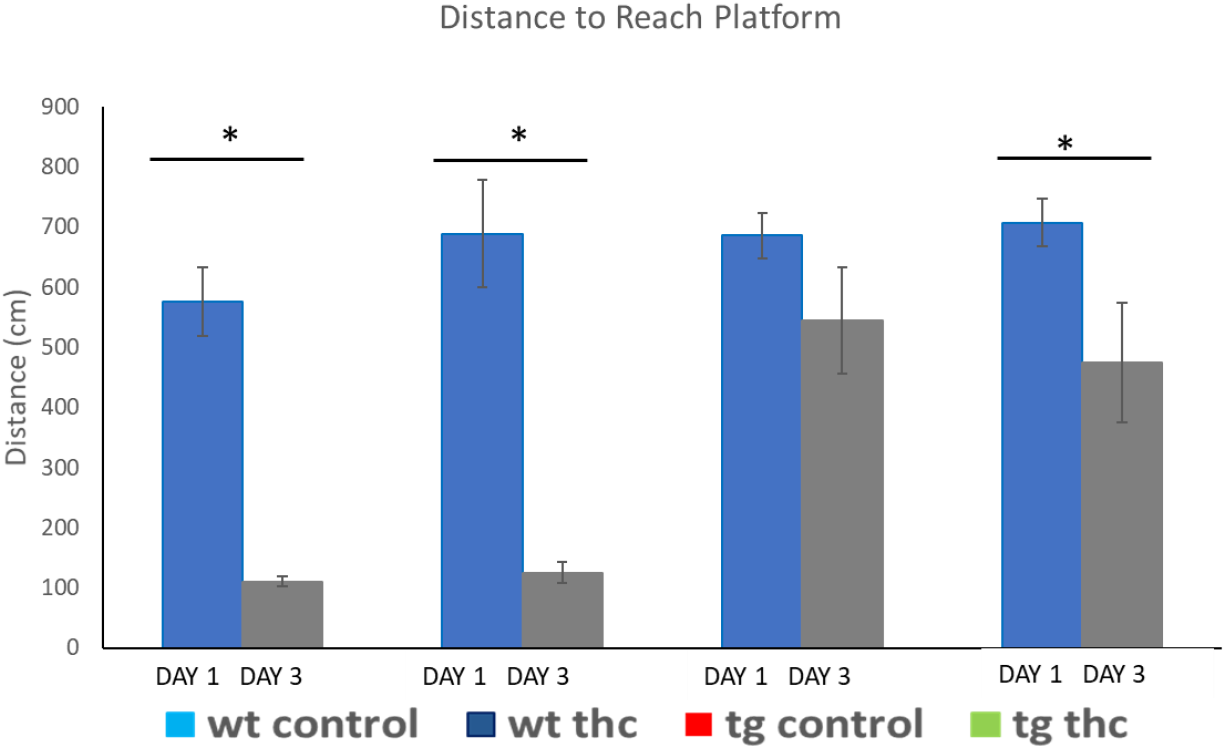
Morris Water Maze. Distance to reach platform. 1-year-old 5xFad and WT littermates were treated with a single I.P. injection of ULD-THC (or vehicle and tested 3 weeks later. Repeated anova yielded significant interaction. Pairwise comparisons showed that while both WT mice and treated 5xFad mice significantly improved from day 1 to day 3.

There was no significant difference in motor function between all groups (Fig 5).

**Figure 5.**
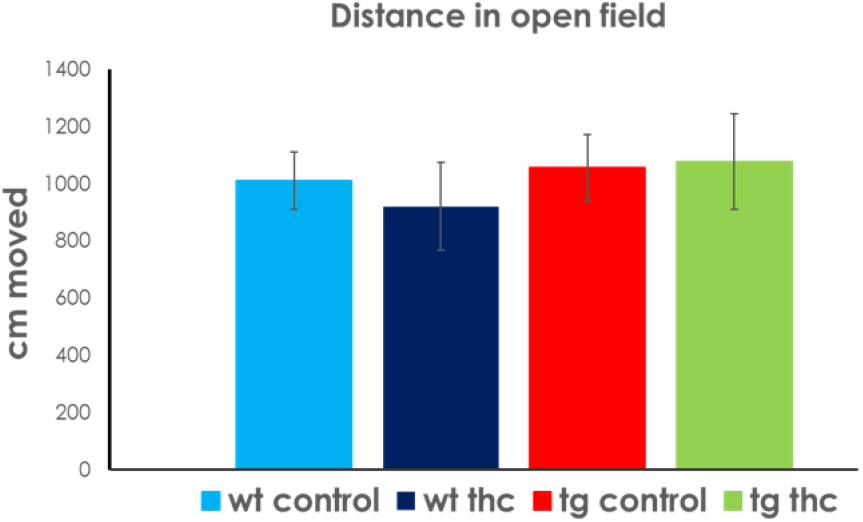
Motor Functions. One-year-old 5xFad (tg) and control (wt) littermates were treated with a single I.P. injection of ULD-THC or vehicle and tested 3 weeks later (N=8 in each group). No significant difference was found between groups in motor function in the open field test.

## Discussion

In this preliminary study, we show for the first time the protective effect of a single treatment of ULD-THC in an AD mice model. 5xFAD mice receiving a single treatment of ULD-THC (0.002 mg/kg) after diseases onset showed improved cognitive performance compared to non-treated AD mice, both in short-term and long-term spatial memory tests.

Our results coincide with previous in-vitro evidence regarding the beneficial effect of THC in AD. CB2R activation in vitro, by a low concentration of a CB2R agonist, resulted in the removal of A_β_ deposits [32]; while in-vivo chronic treatment with CB1R and CB2R agonists improved tau and amyloid pathology in a mouse model of tauopathy [33], and reduced cognitive deficits in mice and rats AD models [34, 35]. Moreover, daily chronic treatment with THC in 5XFAD/APP mice reduced the Aβ burden [36], although it should be noted that no behavioral aspects were examined. In another study, a CB2 receptor agonist affected the oxidative stress response in an APP/PS1 mice model of AD [37], while THC treatment in a mouse model of sporadic AD reduced neuronal loss and restored cerebral glucose metabolism in the hippocampus [38]. All these studies used chronic treatments with conventional (high) doses of cannabinoid agonists (1-20mg/kg). In this regard, one should consider the many limitations of conventional doses of cannabinoids. As mentioned before, numerous studies have shown that THC has deleterious effects [18, 19]. THC exposure in the young has been related to deficits in attention [20], short-term memory [21], and spatial memory [22]. For a more comprehensive review of the possible deleterious effects of THC, see our recent publication [27].

Although our work here did not examine the molecular mechanisms underlying the beneficial effects of ULD-THC, we do have some indications for a possible mechanism. Our previous work showed that the neuroprotective effect of ULD-THC is accompanied by prolonged activation of signaling pathways that are known to mediate neuronal plasticity, survival, and proliferation [27]. Its beneficial effects were mediated by CB1R [25] and accompanied by activation of the ERK/MAPK system in the frontal cortex and the hippocampus; elevation of cAMP response element-binding protein (pCREB) in the hippocampus, and brain-derived neurotrophic factor (BDNF) in the frontal cortex [24]; and upregulation of Sirtuin 1 (Sirt1) in old mice [26].

The Sirtuin family proteins are nicotinamide adenine dinucleotide (NAD)-dependent deacetylases, comprising seven sirtuins in mammals. They play a role in numerous cellular processes that affect longevity, metabolic homeostasis, inflammation alleviation, and health maintenance [39]. Sirt1 plays a vital role in learning and memory in healthy mice [40]; it is also one of the most promising regulators of longevity. The level of Sirt1 was found to decrease during aging [41]. Because Sirt1 regulates aging and metabolic processes, it is also thought to be involved in the pathogenesis of AD [42]. Sirt1 levels are down-regulated in AD patients [43], with a direct connection to Aβ [44] and tau [45] accumulation. Sirt1 activation is thought to play a role in various therapies, which were suggested to improve AD deficits [46]. The involvement of Sirt1 is thought to be related to cAMP responsive-element binding (CREB) [47]. CREB is well known for its metabolic functions [48], but it is also connected to aging and cognitive decline [49]. BDNF, downstream of Sirt1, is also associated with several AD treatments. BDNF is produced by astrocytes [50] and neurons [51] and is a crucial mediator of neuronal plasticity [52] and neurogenesis [53, 54]. BDNF plays an essential role in the development, survival, and maintenance of neurons, has been shown to decrease in late adulthood in correlation with a decline in hippocampal volume [55], and is also connected to age-related cognitive decline [56] and AD [57]. Thus, ULD-THC may upregulate Sirt1 and BDNF expression [24, 26], both related to improved learning and neurogenesis.

Our preliminary data show a long-term effect that can be explained via two possible, although not exclusive, mechanisms– gene regulation and neurogenesis. On the one hand, ULD-THC might induce epigenetic effects by a persistent expression of genes that continuously regulate the production of functional proteins. On the other hand, ULD-THC might lead to the formation of new neurons that either replace dying neurons in the damaged brain or create new neuronal networks[27].

To conclude, the current research demonstrates for the first time the beneficial effects of a single ultra-low dose of THC in a mice model of AD after disease onset.

## Acknowledgements

Funding for this study was provided by the Open University of Israel research fund (http://www.openu.ac.il).

